# ALPHA/BETA HYDROLASE DOMAIN CONTAINING 5 SUPPORTS GLUCOSE-STIMULATED-LIPOLYSIS AND INSULIN SECRETION IN PANCREATIC BETA CELLS

**DOI:** 10.1101/2025.05.21.655350

**Authors:** Lucy Kim, Siming Liu, Spencer Peachee, Syreine Richtsmeier, Claudia Jun, Gianna Sagona, Anamika Vikram, Aidan Freshly, Yumi Imai

## Abstract

PNPLA2 (aka adipose triglyceride lipase) is the rate limiting enzyme of triglyceride (TG) catabolism at the surface of lipid droplets (LDs). Lipolysis by PNPLA2 increases with glucose in pancreatic beta cells and produces metabolites that support glucose-stimulated insulin secretion (GSIS). Upregulation of lipolysis by glucose is blunted in human islets affected by type 2 diabetes. Here, we aim to determine how glucose regulates lipolysis in beta cells.

We found that glucose recruits alpha/beta hydrolase domain containing 5 (ABHD5), a co-activator of PNPLA2 to LDs where lipolysis occurs. ABHD5 recruitment to LDs was reduced by the addition of H-89 and the expression of a mutant ABHD5 that is resistant to phosphorylation by a cAMP dependent kinase (PKA) indicating partial dependence of recruitment on PKA. Importantly, ABHD5 was indispensable for glucose responsiveness of lipolysis in INS-1 832/13 (INS-1) cells. Downregulation of ABHD5 increased LDs and TG content in INS-1 cells and human islets indicating that ABHD5 is highly active in beta cells. Additionally, glucose-stimulated insulin secretion was impaired in ABHD5 downregulated INS-1 and human pseudoislets, which agrees with impaired GSIS in ATGL downregulated beta cells. Thus, ABHD5 plays an important role in conferring glucose responsiveness of lipolysis and supporting insulin secretion in beta cells.

## INTRODUCTION

Beta cell dysfunction is a major culprit for the development and progression of type 2 diabetes (T2D).^1^ One proposed contributor to beta cell dysfunction in T2D is nutrient overload that manifests as the accumulation of lipids within the cells.^1,2^ Such lipids, mainly triglycerides (TGs), are stored in dynamic organelles known as lipid droplets (LDs). LDs consist of a phospholipid monolayer surrounding a core of neutral lipids and allow for the flexible storage and release of core lipids through associated surface proteins and regulators.^3,4^ Patain-like phospholipase domain 2 (PNPLA2, aka as adipose triglyceride lipase, ATGL) catalyzes the breakdown of TGs into diacylglycerides and fatty acids at the LD surface.^5^ Although pancreatic beta cells have a relatively limited number of LDs, PNPLA2-mediated lipolysis is very active and limits TG accumulation in beta cells as indicated by significant accumulation of TGs when PNPLA2 is pharmacologically or genetically suppressed in beta cell lines, mouse beta cells, and human beta cells.^6-9^ More importantly, lipolytic metabolites such as fatty acids and 1-monoacylglyceride are known to support glucose-stimulated insulin secretion (GSIS) by multiple mechanisms including activation of FFAR1 receptor, MUNC13A activation, PPARδ activation, and stabilization of syntaxin 1a (reviewed in^3^). Indeed, pharmacological or genetical suppression of PNPLA2 in beta cells impairs GSIS.^6-9^ Interestingly, beta cell lipolysis is upregulated by glucose, and this process is disrupted in human islets affected by T2D, raising the possibility that impaired lipolysis contributes to impaired GSIS in T2D.^6^ However, how glucose increases lipolysis in pancreatic beta cells is unknown.

The activity of PNPLA2 is known to be regulated by co-factors that positively or negatively regulate its activity.^5^ Hypoxia inducible lipid droplet associated protein (HILPDA)^10^ and G0S2 are examples of inhibitory regulators.^11^ Alpha/beta hydrolase domain containing 5 (ABHD5) is a co-factor that is required for full activity of PNPLA2 lipase activity in liver, muscles, and adipose tissue.^12,13^ Within adipocytes and myocytes, the interaction between PNPLA2 and ABHD5 is regulated through lipid droplet protein perilipin 1 (PLIN1) and PLIN5, respectively.^14,15^ PLIN1 and PLIN5 sequester ABHD5 during the basal condition and release ABHD5 to allow activation of PNPLA2 when PLIN1 and PLIN5 are phosphorylated by cAMP-dependent protein kinase (PKA).^16^ However, the regulation of PNPLA2 in cells that have low expression of PLIN1 and PLIN5 such as pancreatic beta cells is unknown.^17^ PLIN2, a predominant PLIN in beta cells, is not a target of PKA phosphorylation or known to sequestrate ABHD5.^3^

Here, we report that glucose increases the association of ABHD5 with LDs and ABHD5 is required for glucose to stimulate lipolysis in beta cells. Downregulation of ABHD5 in beta cell models increases in TG and LD accumulation significantly and impairs GSIS highlighting that ABHD5 is a key regulator of lipolysis in beta cells.

## RESULTS

### PNPLA2 co-activator ABHD5 is recruited to lipid droplets in response to glucose in beta cells

Although glucose is known to increase lipolysis by PNPLA2 in beta cells,^6^ a molecular mechanism that confers glucose responsiveness of lipolysis in beta cells is unknown. We assessed lipolysis either as the reduction of TG or LD pool when TG synthesis is blocked (Fig. 1).^6,8^ KCl that depolarizes beta cells and triggers insulin secretion is not sufficient to increase lipolysis in INS-1 cells. This points to glucose metabolism upstream of K_ATP_ channel closure as a factor required for glucose to increase lipolysis in beta cells (Fig. 1a). Next, we tested whether PKA contributes to glucose-regulated lipolysis in beta cells considering that glucose is reported to activate PKA in beta cells^18,19^ and PKA is a key regulator of lipolysis in many cells including adipocytes.^5^ A cAMP analog 8Br-cAMP was sufficient to increase lipolysis but not to the extent of 16.8 mM glucose while a PKA inhibitor H-89 partially reduced lipolysis at 16.8 mM glucose (Fig. 1b). Thus, PKA is at least one of the factors that mediate glucose responsiveness of lipolysis in INS-1 cells. PLIN2, a predominant PLIN in beta cells, is not a target of PKA phosphorylation. Considering that ABHD5 positively regulates PNPLA2 and can be phosphorylated by PKA,^20^ we expressed fluorescent protein tagged ABHD5 in INS-1 cells and measured its association with LDs under 2.8 mM and 16.8 mM glucose conditions. Cells exposed to 16.8 mM glucose showed a significant increase in the percentage of LDs surrounded by ABHD5 compared with cells exposed to 2.8 mM glucose (Fig. 1c). Cells exposed to 2.8 mM glucose with 20 μM 8-Br and those exposed to 16.8 mM glucose with 20 μM H-89 showed intermediate ABHD5 recruitment (Fig. 1c), indicating partial dependence of ABHD5 recruitment on PKA that correlates well with partial dependence of lipolysis on PKA. ABHD5 has a PKA consensus sequence at RKYS^239^S^240^ (amino acid numbering based on rat) that is well-conserved among species and phosphorylated by PKA in NIH3T3 cells.^20^ As shown in Fig. 1d-e, expression of a phosphorylation resistant ABHD5 mutant (AA) revealed a partial reduction of recruitment compared with WT ABHD5 at high glucose, corroborating a partial dependence of ABHD5 recruitment on PKA dependent phosphorylation. Expression of the phosphorylation resistant ABHD5 mutant in human beta cells exposed to 16.8 mM glucose conditions demonstrated partial reduction of ABHD5 on the LD surface compared with WT as well (Fig. 1f-g). Collectively, ABHD5 recruitment was indicated as a potential mediator of glucose regulation of lipolysis in beta cells that is partially dependent on phosphorylation by PKA.

**Figure 1.**
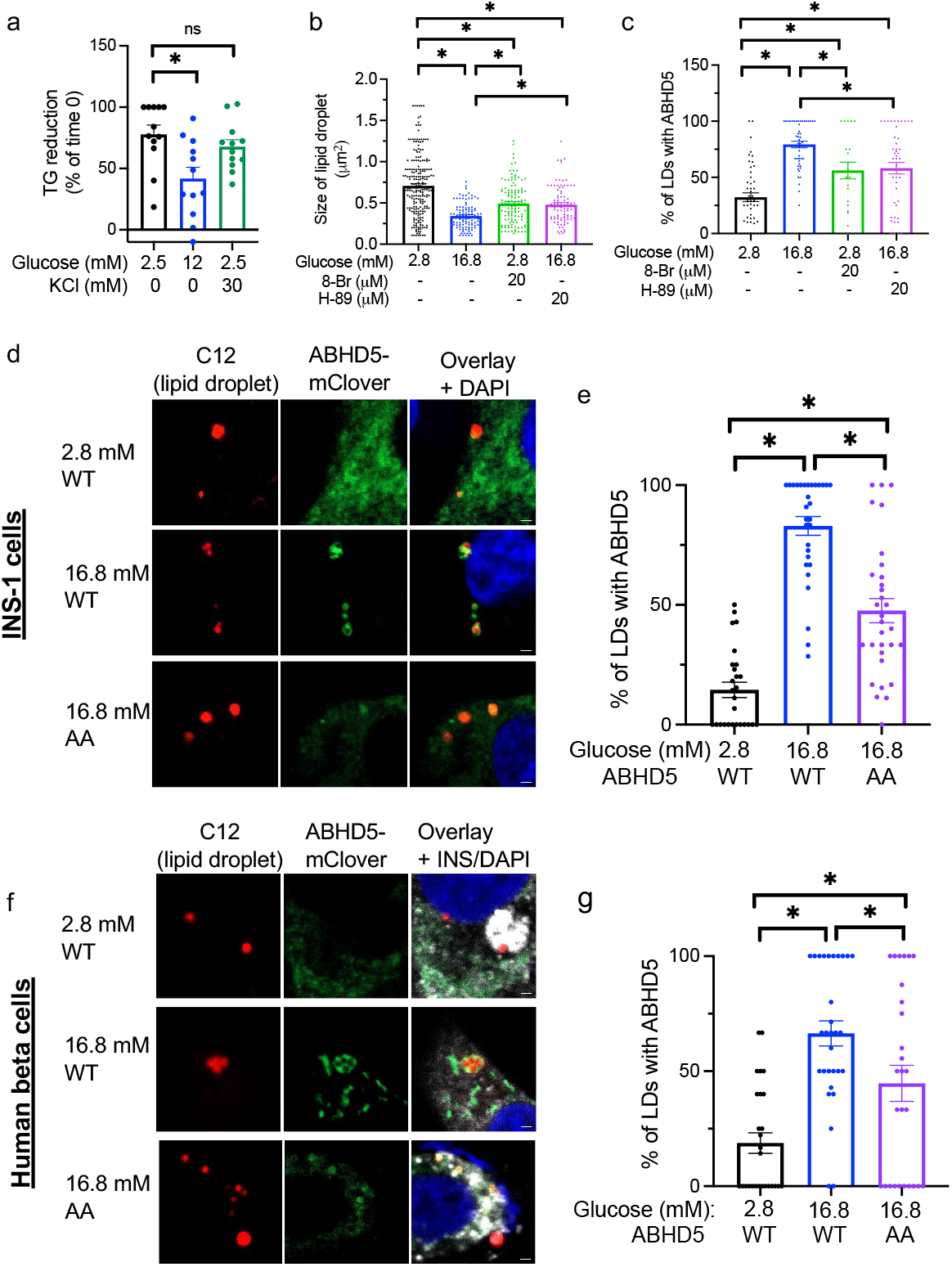
ABHD5 is recruited to lipid droplets at high glucose in beta cells. (a) Lipolysis at 2.5 mM glucose, 12 mM glucose, and 2.5 mM glucose with 30 mM KCl was measured as the reduction of triglycerides (TG) in the presence of a TG synthesis inhibitor triacsin C as in Methods. TG corrected for protein contents were expressed taking TG levels at time 0 as 100%. Data combine five independent experiments. n=12. (b) Size of lipid droplets (LDs) in the presence of triacsin C under indicated concentration of glucose with or without 20 μM 8-Br cAMP (8-Br) or 20 μM H-89. Each dot represents a single LD. n= 89-223. (c) ABHD5-mClover was expressed in INS-1 cells and the proportion of LDs with ABHD5-mClover was measured under indicated concentration of glucose with or without 20 μM 8-Br or 20 μM H-89. Each dot represents a single image. n=20-49. (d-g) WT or PKA resistant mutant (AA) ABHD5-mClover was expressed by lentivirus in INS-1 cells (d, e) or human beta cells (f, g) that were incubated by BODIPY C12 (red) to visualize LD. (d) Representative confocal microscopy images and (e) proportion of LDs with ABHD5-mClover measured under indicated concentration of glucose in INS-1 cells. Each dot represents one image and data combine three independent experiments. n=30-32. (f) Representative confocal microscopy images and (g) proportion of LDs with ABHD5-mClover measured under indicated concentration of glucose in human beta cells visualized by anti-insulin (INS) antibody (white). Data combined images from three independent donors. Scale bar 1 μm. Nuclei were stained with DAPI (blue). Bar graphs show mean ± sem. One way ANOVA with Dunnett’s multiple comparison test (a-c) or Tukey’s multiple comparison test (d, e, g). *; p<0.05.

### ABHD5 is required for glucose responsiveness of lipolysis and limits lipid droplets size in beta cells cultured in a glucose sufficient condition

Next, to confirm that ABHD5 is required for glucose-stimulated lipolysis in INS-1 cells, we downregulated ABHD5 in INS-1 cells using two siRNA that target separate regions of the gene (Fig. 2a-b, supplementary Fig. 1). As shown in Fig 2c, glucose-stimulated lipolysis was blunted in ABHD5 downregulated cells indicating that ABHD5 is necessary for glucose to increase lipolysis in INS-1 cells further supporting that ABHD5 is a key element of glucose-responsive lipolysis in beta cells. The downregulation of ABHD5 markedly increased the size of LDs in INS-1 cells cultured at glucose sufficient condition (11. 1. mM glucose, Fig. 2d-e). The number of LDs was significantly increased in cells treated with siABHD5a and siABHD5b individually compared with cont INS-1 cells (Fig. 2f). Total LD area per cells was markedly increased after downregulation of ABHD5 using any combination of siRNA in INS-1 cells (Fig. 2g).

**Figure 2.**
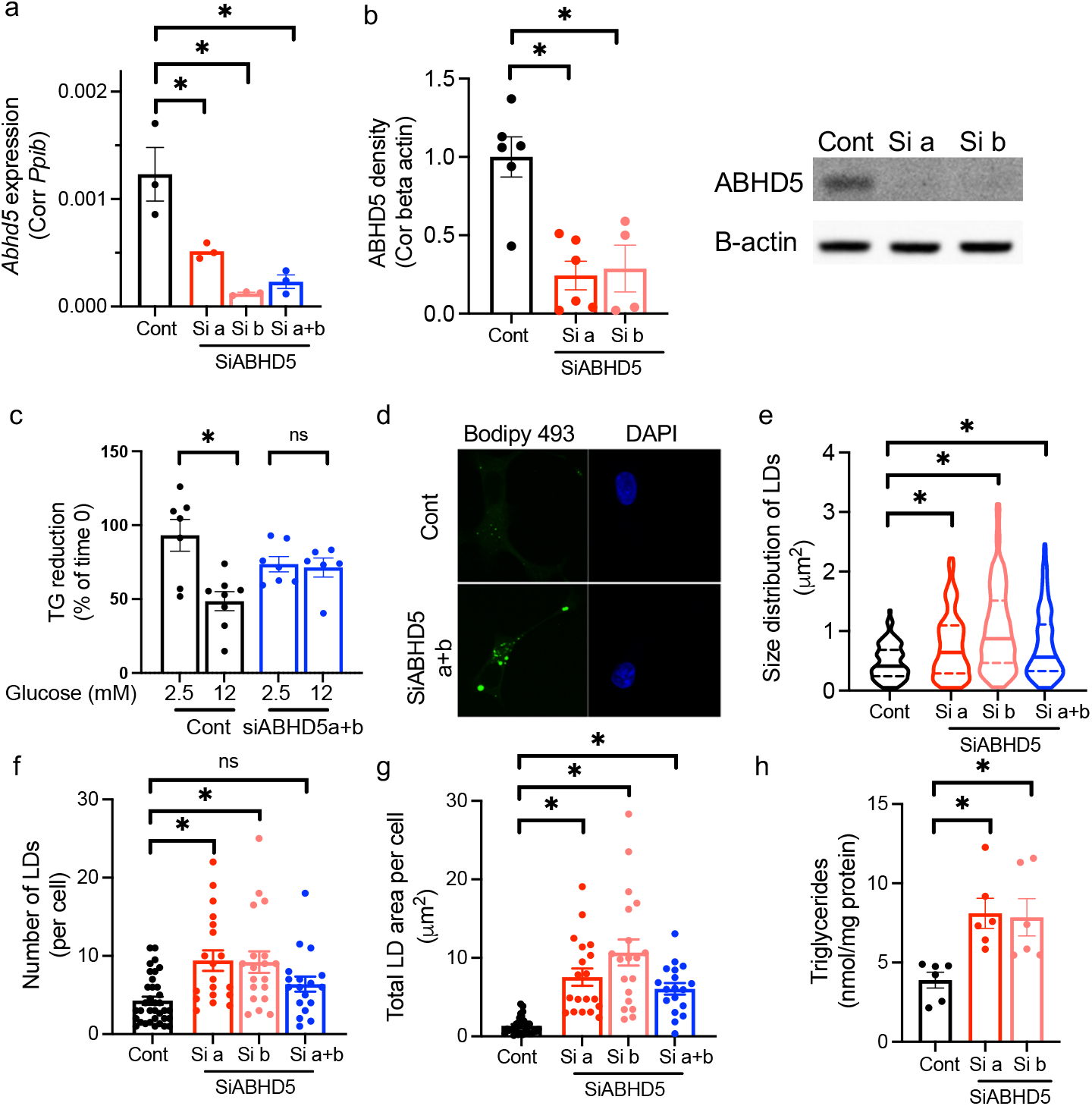
ABHD5 is promoting lipolysis under glucose sufficient condition in INS-1 cells. INS-1 cells were transfected with non-targeting SiRNA (Cont) and two SiRNA against ABHD5 individually (Si a and Si b) or in combination (Si a+b). (a) qPCR assessed *Abhd5*expression using *Ppib* as a housekeeping gene. n=3. Representative of three experiments. (b) Western blot of ABHD5 and beta-actin (B-actin). Band intensity of ABHD5 was corrected for B-actin and expressed taking average value for Cont in each experiment as 1. n= 4-6. A representative image is shown on the right. (c) Lipolysis at 2.5 mM and 12 mM glucose was measured as the reduction of TG in the presence of a triglycerides (TG) synthesis inhibitor triacsin C. Data combine three independent experiments. n=8-6. (d-g) Lipid droplets (LDs) in INS-1 cells were visualized by BODIPY 493/503 (green) under confocal microscopy and analyzed for size and number. Nuclei were stained with DAPI (blue). (d) Representative images. (e) Violin plots of LD size distribution. Median and quantiles are indicated. n= 203 to 226 LDs. (f) The number of LDs per cell. Each dot represents a single cell. n=18 to 35 cells and (g) the sum of LD covered area in each cell. Each dot represents a single cell. n= 18 to 31 cells. (d-g) Combined data from four independent experiments. (g) TG contents corrected for protein contents. Data combine results of three independent experiments each performed in duplicates. n = 6. Bar graphs show mean ± sem. One way ANOVA with Dunnett’s multiple comparison test (a-g) or Student’s t test (h) were performed. ns; not significantly different. *; p<0.05.

In concordance with the increase in total LD area per cell, TG, the major constituent of LDs in beta cells, was increased two-fold in ABHD5 downregulated INS-1 cells (Fig. 2h). Data indicate the magnitude by which ABHD5 contributes to the regulation of lipolysis and LD pool size is significant in beta cells.

We next determined whether human beta cells also increase LDs when ABHD5 is downregulated (Fig. 3a-b). Pairwise comparison showed 1.79 times increase in average size of LDs and 2.02 increase in the number of LDs for lenti-shABHD5-treated beta cells compared with lenti-scramble shRNA treated cells (Fig. 3c-e), which is in alignment with the increased size and number of LDs in siABHD5 treated INS-1 data (Fig 2d-g). As expected from the increase in size and number of LDs, total LD area corrected for insulin positive area in shABHD5 islets was 3.85 times higher than control beta cells (Fig. 3f). TG contents were also increased in human pseudoislets after ABHD5 downregulation (Fig. 3g). Thus, both INS-1 cells and human islets showed increase in LDs and TG contents after ABHD5 downregulation, indicating that ABHD5 is critical for lipolysis in beta cells under glucose sufficient condition and limits LD accumulation in beta cells.

**Figure 3.**
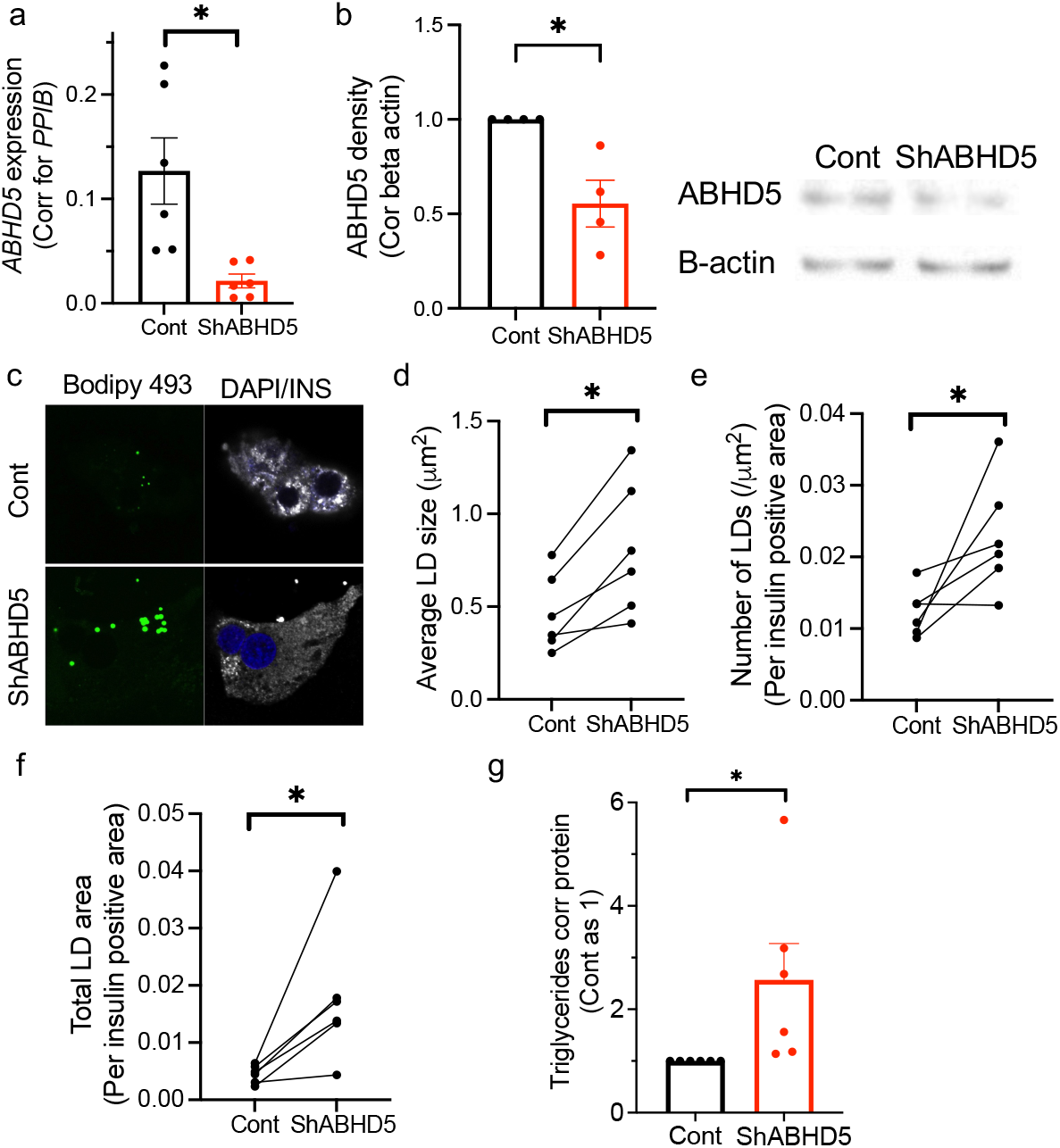
ABHD5 is promoting lipolysis under glucose sufficient condition in human beta cells. (a) qPCR compared the expression of ABHD5 in human pseudoislets transduced with lentivirus carrying scramble RNA (Cont) or shRNA against ABHD5 (shABHD5). Expression was normalized to *PPIB*. n= 6 donors. (b) Western blot of ABHD5 and beta-actin (B-actin). Band intensity of ABHD5 was corrected for B-actin and expressed taking average value for Cont in each experiment as 1 from n= 4 donors. A representative image is shown on the right. (c-e) LD size and numbers were determined in human beta cells treated with lenti-shABHD5 by visualizing LDs with BODIPY 493/503 (green) and imaging under confocal microscopy. Beta cells were identified by anti-insulin (INS) antibody (white) and nuclei by DAPI (blue). (c) Representative images. (d) Average size of LDs in beta cells, (e) number of LD corrected for insulin positive area, and (f) the sum of LD covered area corrected for insulin positive area for n= 6 donors. Each dot represents an average value obtained from a single donor and data from the same donor is connected by a line. (g) Triglycerides (TG) content corrected for protein content were compared between human pseudoislets treated with cont and lenti-shABHD5. Value for Cont was taken as 1 for each donor. N = 6 donors. Data are mean ± sem. * p<0.05 by Student’s t test.

### ABHD5 downregulation impairs insulin secretion in INS-1 cells and human pseudoislets

Downregulation of PNPLA2 impairs glucose-stimulated insulin secretion (GSIS) in INS-1 cells and human islets indicating important contribution of lipolytic metabolites for GSIS.^6,7,21^ If ABHD5 actively regulates lipolysis in beta cells, we expect that downregulation of ABHD5 will impair GSIS as for PNPLA2 downregulation. Thus, we then assessed GSIS after ABHD5 downregulation. While high glucose increased insulin secretion in both control and siABHD5 treated INS-1 cells, the increase in insulin secretion in response to high glucose was reduced to 63.0% of control cells indicating GSIS is blunted in ABHD5-deficient cells (Fig. 4a). Total insulin content corrected for protein was similar across all treatment groups (Fig. 4b).

**Figure 4.**
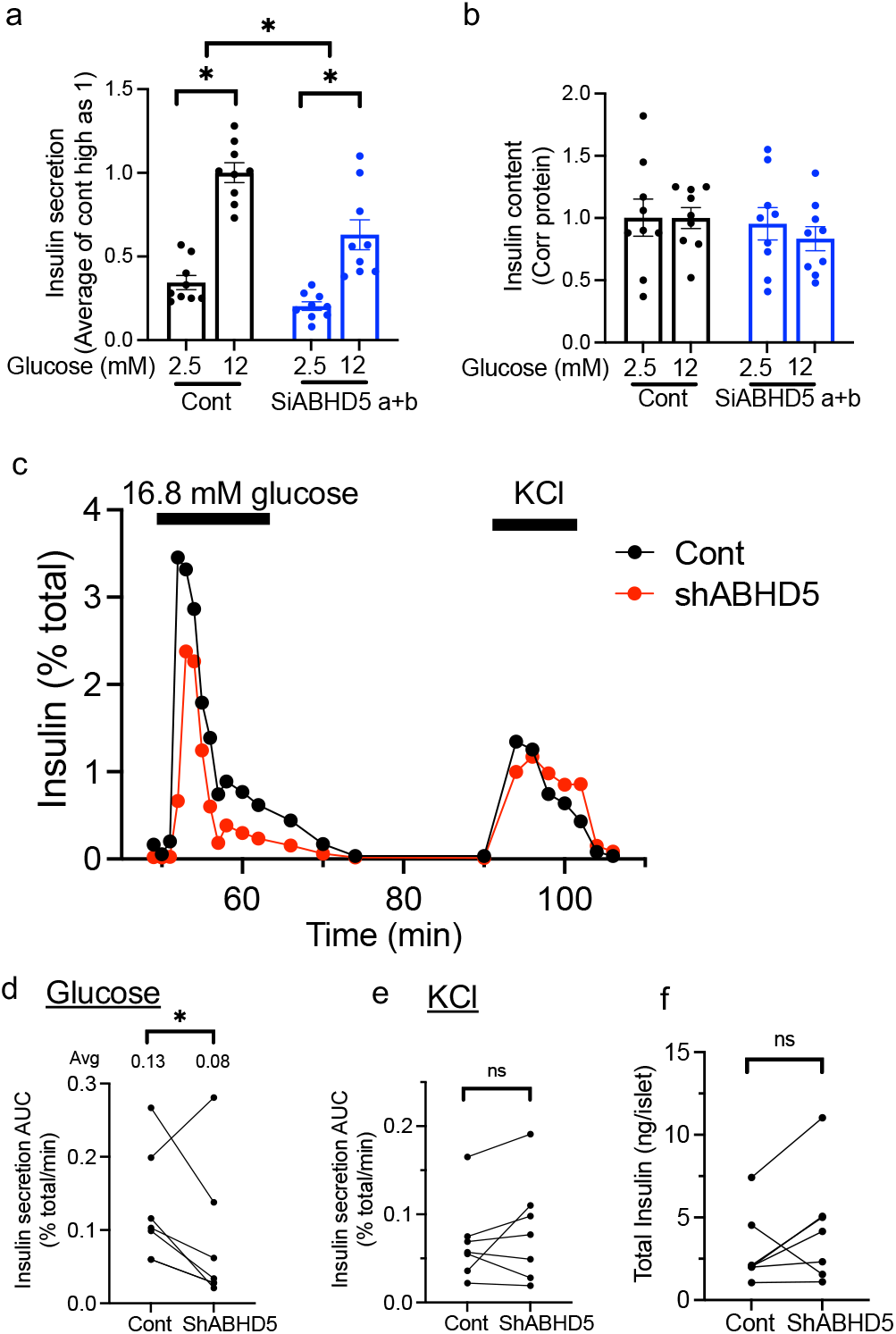
Downregulation of ABHD5 impairs glucose-stimulated insulin secretion from beta cells. (a-b) Insulin secretion at the indicated concentrations of glucose was compared between INS-1 cells treated with control SiRNA (Cont) or mixture of two siRNA against ABHD5 (Si a+b). n=9 combining data from 3 experiments, each performed in triplicates. (a) Secreted insulin was corrected for insulin content and expressed taking an average value for high glucose in Cont as 1. (b) Insulin content in cell lysate was corrected for protein content and expressed taking an average value for high glucose in cont as 1. Data are mean ± sem. *; p<0.05 by One way ANOVA with Sidak’s multiple comparison test. (c-f) Insulin secretion assessed by perifusion in human pseudoislets transduced by lenti-scr (Cont) and lenti-shABHD5 (shABHD5). Insulin secretion was corrected for total insulin contents. (c) A representative insulin secretion profile in response to 16.8 mM glucose (G) and 30 mM KCl at times indicated by bars. (d) Area under the curve (AUC) of insulin secretion under 16.8 mM glucose, (e) AUC of insulin secretion under 30 mM KCl, and (f) total insulin content. Each dot represents a single donor and data from the same donor are connected by a line. n = 6 donors. * p<0.05 by paired Student’s t test.

We also tested insulin secretion by perifusion of human pseudoislets in which ABHD5 was downregulated using lenti-shRNA (Fig. 3a, b). GSIS was significantly blunted in human pseudoislets after ABHD5 downregulation with less effect on KCl stimulated insulin secretion (Fig. 4c-e). There was no significant difference in total insulin contents (Fig. 4f).

## DISCUSSION

Previous studies have shown that lipolysis by PNPLA2 supports insulin secretion in beta cells.^6,7,21^ Interestingly, glucose responsiveness of lipolysis is blunted in human islets from T2D donors and potentially contributes to impaired beta cell function in T2D.^6^ Thus, the current study aimed to determine a factor that mediates the increase of lipolysis by glucose in beta cells. We found that ABHD5, a PNPLA2 activator, is recruited to LDs by glucose, a process that is partly dependent on phosphorylation of ABHD5 by PKA. Downregulation of ABHD5 blunts the increase of lipolysis by glucose in INS-1 cells providing additional support that ABHD5 is critical to confer glucose responsiveness to lipolysis in beta cells. Substantial accumulation of TG and LDs at glucose sufficient conditions in ABHD5 deficient cells indicates that lipolysis is highly active in beta cells and ABHD5 significantly supports lipolysis in beta cells. As expected from the role of lipolysis in supporting GSIS, ABHD5 downregulation impairs GSIS in ABHD5-deficient INS-1 cells and human pseudoislets. Thus, our results show the significant role played by ABHD5 to activate PNPLA2 and aid in cell function in pancreatic beta cells.

In beta cells, the association of ABHD5 with LDs is increased by glucose when lipolysis is active. Considering that PNPLA2 acts on the surface of LDs to catalyze TG degradation,^13^ the recruitment of ABHD5 to LDs at a glucose-stimulated condition appears to be logical. However, this contrasts with ABHD5 distribution at LDs in non-stimulated basal conditions in cells that express perilipin 1 (PLIN1) and PLIN5. PLIN1 is highly expressed in adipocytes and PLIN5 in oxidative cells such as skeletal muscle, cardiomyocytes, and brown adipocytes.^16^ In these cells, ABHD5 is bound to PLIN1 and PLIN5 that are anchored at the LD surface at the basal condition to sequester ABHD5 away from PNPLA2. Upon phosphorylation of PLIN1 and PLIN5 by PKA, ABHD5 is released from PLINs to bind PNPLA2 to increase lipolysis.^14,15^ Since the expression of PLIN1 and PLIN5 is very low in beta cells under a nutrient sufficient condition, PLIN1 and PLIN5 are unlikely to regulate glucose-responsiveness of lipolysis in beta cells.^17^ Instead of sequestered from PNPLA2 by PLIN1 and PLIN5 on the LD surface at the basal condition,^14,15,20^ ABHD5 in beta cells appears to be dissociated from LD surface at the basal condition when glucose level is low and increases the association with LDs upon glucose stimulation to activate PNPLA2 at the surface of LDs. Therefore, the regulation of ABHD5 localization in cells with low expression of PLIN1 and PLIN5 is distinct from those with high expression of PLIN1 and PLIN5. It remains to be determined how glucose changes the location of ABHD5 in beta cells. A previous study showed that PKA dependent phosphorylation of ABHD5 does not alter its potency as a co-activator of PNPLA2 but is required for dissociation from PLIN1 coated LDs in adipocytes upon stimulation.^20^ The partial reduction of ABHD5 association with LDs at high glucose in the presence of H-89 or phosphorylation resistant mutant indicates that PKA dependent phosphorylation of ABHD5 facilitates glucose-stimulated ABHD5 association with LDs in beta cells at least partly. However, a PKA independent factor also appears to contribute to ABHD5 recruitment to LDs at high glucose conditions, indicating multimodal regulation of ABHD5 distribution by glucose in beta cells.

Significant accumulation of TG and LDs in ABHD5 deficient beta cells indicates ABHD5 avidly supports lipolysis by PNPLA2 in glucose sufficient conditions. Thus, the reduction in GSIS in ABHD5 deficient beta cells is expected considering that ATGL deficiency in beta cells impairs GSIS.^6,7,21^ However, it is of note that ABHD5 knockdown or mutations does not phenocopy that of PNPLA2 perfectly in other cells, which is partly due to the interaction of ABHD5 with proteins besides PNPLA2.^22^ In keratinocytes, ABHD5 interacts with PNPLA1 and the interaction between the two is considered to regulate omega-O-acyl ceramide production that is essential for skin barrier formation.^23^ Ichthyosis, one of phenotypes seen in ABHD5 mutations in humans is considered to be due to the impaired interaction of ABHD5 with PNPLA1 and not seen in human with PNPLA2 mutations.^22^ PNPLA3 is highly expressed in hepatocytes and its PNPLAD3 I148M variant increases a risk of metabolic dysfunction-associated fatty liver disease in humans. In hepatocytes, the competition for ABHD5 between PNPLA2 and PNPLA3 is proposed to provide an additional layer of regulation of lipolysis by PNPLA2.^24^ The expression of PNPLA1 and PNPLA3 is very low in beta cells based on single cell sequence data.^25,26^ However, it is plausible there are additional targets of ABHD5 in beta cells.

As above, the mutation of ABHD5 in humans causes Chanarin-Dorfman syndrome, neutral lipid storage disease with ichthyosis.^12^ While hepatosteatosis and other factors may contribute, the development of T2D is reported in Chanarin-Dorfman syndrome,^27,28^ which does not contradict the beta cell dysfunction demonstrated in ABHD5 deficient beta cell models. Beside a monogenic form of ABHD5 mutation, the functional impairment of ABHD5 recruitment to LDs may occur in T2D when glucose metabolism is dysregulated.^1^ However, the impact of ABHD5 deficiency in beta cells in vivo needs further confirmation. We have not been able to use a mouse model of ABHD5 knockout as LDs are substantially smaller in mouse beta cells and glucose responsiveness of lipolysis was not clear in mouse islets.^6,17^ It also needs to be noted that PNPLA2 is also regulated by inhibitory cofactors such as HILPDA.^10^ It remains possible that upregulated HILPDA activity may account for dysregulated lipolysis and beta cell function in addition to ABHD5 dysfunction in T2D islets.

In summary, our study demonstrated ABHD5 recruitment to LDs as one mechanism to increase lipolysis in response to glucose. ABHD5 is required for glucose responsiveness of lipolysis in beta cells and contributes to the regulation of insulin secretion.

## MATERIALS AND METHODS

### INS-1 cells

823/13 cells (INS-1 cells, a gift from Dr. Christopher Newgard (Duke University, Durham NC)) were cultured in INS-1 medium comprised of RPMI1640 supplemented with 10 mM HEPES, 10% FBS, 2 mM L-glutamine, 1 mM sodium pyruvate, and penicillin+ streptomycin (Pen-Strept). To downregulate ABHD5, cells were transfected with 30 nM of siRNA targeting ABHD5 (siABHD5, rn.Ri.Abhd5.13.2 (si a) and/or rn.Ri.Abhd5.13.3 (si b) from Integrated DNA Technologies (IDT)) using DharmaFECT1 transfection reagent as published.^6^ Non-targeting siRNA (IDT) served as negative control. Subsequent experiments were performed 72 h after transfection. For recruitment imaging experiments, INS-1 cells were transduced by lentivirus expressing rat ABHD5-mClover and/or rat ABHD5-mClover with S239A and S240A mutation (AA) and cells were analyzed 72 h after transduction.

### Human islets

Human islets from Integrated Islet Distribution Program (IIDP), Alberta Diabetes Institutes, or Prodo labs (Table 1 for donor characteristics) with reported viability and purity above 80% were cultured overnight at 37 ºC and 5% CO_2_ upon arrival for recovery from shipping. Table 1 used data from the Organ Procurement and Transplantation Network (OPTN). The OPTN data system includes data on all donor, wait-listed candidates, and transplant recipients in the US, submitted by the members of the Organ Procurement and Transplantation Network (OPTN). The Health Resources and Services Administration (HRSA), U.S. Department of Health and Human Services provides oversight to the activities of the OPTN contractor.

**Table 1.** Characteristics of human islet donors.

| Donor Age<br>(years) | Donor Sex<br>(M/F) | Donor BMI<br>(kg/m<sup>2</sup>) | Donor<br>HbA1c | Origin/<br>source of islets<sup>#</sup> | Data generated |
| --- | --- | --- | --- | --- | --- |
| 54 | M | 30.8 | 5.1 | IIDP | Fig. 3a, 4d-f |
| 38 | M | 28.4 | 6.1 | IIDP | Fig. 3a, g, 4d -f |
| 30 | F | 39.9 | 5.3 | IIDP | Fig. 3a |
| 34 | M | 25.8 | 5.0 | IIDP | Fig. 3a, Fig. 4c-f |
| 16 | M | 29.6 | 5.4 | IIDP | Fig. 3d-f |
| 28 | M | 29.3 | 5.0 | IIDP | Fig. 3a, g |
| 20 | M | 38.3 | 5.5 | IIDP | Fig. 3g, 4d-f |
| 55 | F | 31.7 | 5.9 | IIDP | Fig. 3g, 4d-f |
| 61 | M | 24.6 | 5.4 | IIDP | Fig. 3g, 4d-f |
| 21 | M | 38.1 | 5.0 | IIDP | Fig. 3b |
| 39 | M | 27.1 | 5.3 | IIDP | Fig. 3b |
| 33 | M | 28.7 | 5.5 | IIDP | Fig. 3b |
| 56 | M | 29.3 | 5.4 | IIDP | Fig. 1f |
| 39 | M | 22.9 | 5.7 | IIDP | Fig. 1f |
| 66 | M | 29.6 | 5.4 | ADI | Fig. 3a, d-g, 4d-f |
| 62 | M | 30.1 | 4.9 | ADI | Fig. 3d-f |
| 34 | M | 24.2 | 5.1 | ADI | Fig. 1f |
| 36 | M | 24.4 | 5.7 | Prodo lab | Fig. 3d-f |
| 36 | M | 34 | 4.9 | Prodo lab | Fig. 3d-f |

<sup>#</sup>: IIDP-Integrated Islet Distribution Program, ADI- Alberta Diabetes Institute

Lentivirus carrying scramble (CCTAAGGTTAAGTCGCCCTCG) and shRNA sequences targeting human ABHD5 (CCG GCC TGA TTT CAA ACG AAA GTA TCT CGA GAT ACT TTC GTT TGA AAT CAG GTT TTT G and CCGGCCTAC GTC TAT ATC ACA CCT TCT CGA GAA GGT GTG ATA TAG ACG TAG GTT TTT) were obtained from the Genetic Perturbation Platform (https://portals.broadinstitute.org/gpp/public). Pseudoislets transduced with lentivirus were created as published.^29^ The study was determined to be a non-human study by Institutional Review Board at University of Iowa.

### Morphometry of LDs and ABHD5 Recruitment in INS-1 cells and human beta cells

INS-1 cells were plated onto a cover glass coated with extracellular matrix of HTB-9 cell as published.^30^ Human islet cells were grown on a confocal dish coated with type IV collagen (C5533 from Sigma) as published.^31^ Cells were fixed for 15 min at 37°C in 4% paraformaldehyde and incubated with 2 μM Bodipy 493/503 (Bodipy 493, Molecular Probes) and 1 μg/ml of DAPI as published.^32^ Islets were also immunostained for anti-insulin (INS, rabbit anti-INS1 antibody, AB_3673135, C27C9, Cell signaling at 1:300) to visualize beta cells. For localization of ABHD5-mClover, cells were incubated in medium containing 3 μM Bodipy C12 (Molecular Probes) with 0.1 mM oleic acid to visualize LDs at 37 °C at 5% CO_2_ for the last 16 h of culture, then incubated in media containing 20 μM Triacsin C in addition to the stated conditions (2.8 mM glucose or 16.8 mM glucose with or without 20 μM H 89 dihydrochloride (H-89, Cayman Chemical, CAS 130964-39-5) or 20 μM 8-bromo Cyclic AMP (8-Br, Cayman Chemical, CAS 76939-46-3)) for 90 min prior to fixation. Images of the cells were captured by a Zeiss 980 microscope. Size and number of LDs were measured in each image using the Bodipy 493 or Bodipy C12 channel in the ImageJ software as published.^31^ To measure ABHD5 recruitment to LD surface, C12-labeled LDs and ABHD5 signals were independently thresholded and added as separate regions of interest (ROI) in ImageJ. Each independent ROI and the overlap of ROIs was recorded and the visualized LD ROIs with surrounding ABHD5 ROI was counted. The same data extraction was repeated for every image in each dataset with the number of LDs with ABHD5 being expressed as a part of the total LDs in each image.

### Measurement of TG contents and Lipolysis Assay

INS-1 cell lysates and islet lysates were prepared in RIPA (Radio-Immunoprecipitation Assay) buffer (Sigma-Aldrich) with 0.01 volumes of Protease Inhibitor Cocktail (P8340, Sigma Aldrich). Protein concentration of the cell lysate was measured using RC DC protein assay kit (Bio-Rad) and Pierce BCA protein assay kit (Thermo fisher). Then, TGs were extracted from the cell lysate by Folch buffer and subsequently measured by Infinity Reagents (Fisher Scientific).^6^ To measure lipolysis as reduction of TG, cells and islets were incubated in the presence of 20 μM triacsin C to block TG synthesis and 5 μM lalistat 2 (LIPA inhibitor) to block lipophagy following a previously published protocol.^6^ In brief, INS-1 cells preincubated in Krebs-Ringer (KRB) buffer with 2 mM glucose, Triacsin C, and lalistat 2 for 60 min was harvested as time 0 samples. Then, cells were incubated with 2 mM glucose, 12 mM glucose, or 30 mM KCl plus Triacsin C for 120 min. TG contents corrected for proteins were expressed taking the value at time 0 as 100%.

### Quantitative PCR

RNA was isolated from INS-1 cells using Invitrogen PureLink RNA Mini Kit (ThermoFischer Scientific) and from islets using TRIZOL reagent (Thermo Fisher), and subsequently reverse transcribed into cDNA as published.^6^ Gene expression was assessed using ABI TaqMan commercial primers (Applied Biosystems).

### Immunoblotting

INS-1 cells and human pseudoislets were solubilized in RIPA with 0.01 volumes of Protease Inhibitor Cocktail (P8340, Sigma Aldrich). 10 to 20 μg/lane of protein were loaded for SDS-PAGE and Western blot as published.^6^ After transfer, nitrocellulose membranes were blocked in 3-5% milk in 1XPBST (PBS containing 0.1% Tween 20) and incubated with primary antibodies followed by secondary antibodies. Membranes were visualized by the SuperSignal West Pico with SuperSignal West Atto (1:40 dilution) chemiluminescent detection system as published^32^ or fluorescence with the iBright FL1500 Imaging System. Densitometric analysis was conducted using ImageJ software (ImageJ.nih.gov). Primary antibodies used were mouse anti-ABHD5 (sc-376931, Santa Cruz Biotechnology, 1:200) and mouse anti-beta-Actin (sc-47778, Santa Cruz Biotechnology, 1:5000). Secondary antibodies were mouse IgG kappa binding protein (m-IgGκ BP)-HRP (sc-2357, Santa Cruz Biotechnology, 1:3000) and IRDye 800CW Goat anti-Mouse IgG at 1:10,000.

### Glucose-stimulated insulin secretion

INS-1 cells were first incubated with 0.25% BSA Perifusion Buffer (120 mM NaCl, 4.8 mM KCl, 2.5 mM CaCl_2_ 2H_2_O, 1.2 mM MgCl_2_ 6H_2_O2.5 mM, 10 mM HEPES, 24 mM NaHCO_3_) containing 2.5 mM glucose at 37°C, 5% CO_2_ for 0.5 hours x twice. Then, cells receive either 2.5 mM glucose Perifusion Buffer or 12 mM glucose Perifusion Buffer and incubated for 2 h. At the end of the incubation period, medium and cell lysates were harvested. Insulin secreted and insulin contents were measured with STELLUX Chemiluminescent rodent Insulin ELISA (ALPCO). Human pseudoislets were perifused by BioRep Perifusion System (BioRep) as published.^29^ In brief, islets were perifused by 0.25% BSA Perifusion Buffer with 2.8 mM glucose for 48 minutes followed by perifusion Buffer containing 16.8 mM glucose or 30 mM KCl plus 2.8 mM glucose as indicated in Fig. 4c. Total insulin contents were obtained from islets incubated overnight at 4 °C in RIPA buffer containing protease inhibitors. Human insulin was measured using STELLUX Chemiluminescent Human Insulin ELISA.

### Statistics

Data are presented as mean ± sem unless otherwise stated in the figure legends. Differences of numeric parameters between two groups were assessed with Student’s t-tests. Multiple group comparisons used one-way ANOVA with a post hoc test as indicated. Outlier assessment was performed by ROUT test. All statistical analyses used Prism 10 (GraphPad, La Jolla, CA). A p < 0.05 was considered significant.

## Data and Resource Availability

The datasets generated during and/or analyzed during the current study are available from the corresponding author upon reasonable request.

## Author Contribution

YI conceived the study and is responsible for all contents of the manuscript. LK (all aspects), SL, (imaging and human islet studies), SP (cell culture, lipolysis assay), SR (ELISA), CJ (Western blot, GS (qPCR), AV (human islet studies), and AF (imaging), were responsible for the acquisition and/or analysis of the data. YI, LK, and SL designed research, drafted the manuscript, and critically revised the manuscript for important intellectual content. All authors revised and approved the final version of the manuscript.

## Acknowledgement

This work was financially supported by National Institutes of Health to YI (R01-DK090490), and VA merit award to YI (I01 BX005107). YI is supported by the Fraternal Order of Eagles Diabetes Research Center. A part of human pancreatic islets was provided by the NIDDK-funded Integrated Islet Distribution Program (IIDP) (RRID:SCR _014387) at City of Hope, NIH Grant # 2UC4DK098085. Human islets were also provided by the Alberta Diabetes Institute Islet Core at the University of Alberta in Edmonton with the assistance of the Human Organ Procurement and Exchange (HOPE) program, Trillium Gift of Life Network (TGLN), and other Canadian organ procurement organizations. A Zeiss 980 confocal microscope located in the University of Iowa Central Microscopy Research Facility (CMRF) was funded by the Roy J Carver Charitable Trust. The data reported here have been supplied by UNOS as the contractor for the Organ Procurement and Transplantation Network (OPTN). The interpretation and reporting of these data are the responsibility of the authors and in no way should be seen as an official policy of or interpretation by the OPTN or the U.S. Government.

## References

1. Imai Y E.-L.D.,, Peachee S J. . Pancreatic islet adaptation and failure in metabolic syndrome. in Metabolic Syndrome, Comprehensive Textbook (ed. AhimA, R.) 385–404 (Springer Nature, 2024).

2. Horii, T., et al. Lipid droplet accumulation in beta cells in patients with type 2 diabetes is associated with insulin resistance, hyperglycemia and beta cell dysfunction involving decreased insulin granules. Front Endocrinol (Lausanne) 13, 996716 (2022).

3. Tong, X., Liu, S., Stein, R. & Imai, Y. Lipid Droplets’ Role in the Regulation of beta-Cell Function and beta-Cell Demise in Type 2 Diabetes. Endocrinology 163, bqac007 (2022).

4. Mathiowetz, A.J. & Olzmann, J.A. Lipid droplets and cellular lipid flux. Nat Cell Biol 26, 331–345 (2024).

5. Grabner, G.F., Xie, H., Schweiger, M. & Zechner, R. Lipolysis: cellular mechanisms for lipid mobilization from fat stores. Nat Metab 3, 1445–1465 (2021).

6. Liu, S., et al. Adipose Triglyceride Lipase Is a Key Lipase for the Mobilization of Lipid Droplets in Human beta-Cells and Critical for the Maintenance of Syntaxin 1a Levels in beta-Cells. Diabetes 69, 1178–1192 (2020).

7. Tang, T., et al. Desnutrin/ATGL activates PPARdelta to promote mitochondrial function for insulin secretion in islet beta cells. Cell Metab 18, 883–895 (2013).

8. Kim, L.B., et al. Acute Inhibition of Adipose Triglyceride Lipase by NG497 Dysregulates Insulin and Glucagon Secretion from Human Islets. Endocrinology (2025).

9. Peyot, M.L., et al. Adipose triglyceride lipase is implicated in fuel- and non-fuel-stimulated insulin secretion. J Biol Chem 284, 16848–16859 (2009).

10. Padmanabha Das, K.M., et al. Hypoxia-inducible lipid droplet-associated protein inhibits adipose triglyceride lipase. J Lipid Res 59, 531–541 (2018).

11. Zhang, X., Heckmann, B.L., Campbell, L.E. & Liu, J. G0S2: A small giant controller of lipolysis and adipose-liver fatty acid flux. Biochim Biophys Acta 1862, 1146–1154 (2017).

12. Lass, A., et al. Adipose triglyceride lipase-mediated lipolysis of cellular fat stores is activated by CGI-58 and defective in Chanarin-Dorfman Syndrome. Cell Metab 3, 309–319 (2006).

13. Vieyres, G., Reichert, I., Carpentier, A., Vondran, F.W.R. & Pietschmann, T. The ATGL lipase cooperates with ABHD5 to mobilize lipids for hepatitis C virus assembly. PLoS Pathog 16, e1008554 (2020).

14. Granneman, J.G., Moore, H.P., Krishnamoorthy, R. & Rathod, M. Perilipin controls lipolysis by regulating the interactions of AB-hydrolase containing 5 (Abhd5) and adipose triglyceride lipase (Atgl). J Biol Chem 284, 34538–34544 (2009).

15. Granneman, J.G., Moore, H.P., Mottillo, E.P. & Zhu, Z. Functional interactions between Mldp (LSDP5) and Abhd5 in the control of intracellular lipid accumulation. J Biol Chem 284, 3049–3057 (2009).

16. Sztalryd, C. & Brasaemle, D.L. The perilipin family of lipid droplet proteins: Gatekeepers of intracellular lipolysis. Biochim Biophys Acta Mol Cell Biol Lipids 1862, 1221–1232 (2017).

17. Imai, Y., Cousins, R.S., Liu, S., Phelps, B.M. & Promes, J.A. Connecting pancreatic islet lipid metabolism with insulin secretion and the development of type 2 diabetes. Ann N Y Acad Sci 1461, 53–72 (2020).

18. Nesher, R., et al. Beta-cell protein kinases and the dynamics of the insulin response to glucose. Diabetes 51 Suppl 1, S68–73 (2002).

19. Ramos, L.S., Zippin, J.H., Kamenetsky, M., Buck, J. & Levin, L.R. Glucose and GLP-1 stimulate cAMP production via distinct adenylyl cyclases in INS-1E insulinoma cells. J Gen Physiol 132, 329–338 (2008).

20. Sahu-Osen, A., et al. CGI-58/ABHD5 is phosphorylated on Ser239 by protein kinase A: control of subcellular localization. J Lipid Res 56, 109–121 (2015).

21. Zhao, S., et al. alpha/beta-Hydrolase domain-6-accessible monoacylglycerol controls glucose-stimulated insulin secretion. Cell Metab 19, 993–1007 (2014).

22. Schratter, M., Lass, A. & Radner, F.P.W. ABHD5-A Regulator of Lipid Metabolism Essential for Diverse Cellular Functions. Metabolites 12(2022).

23. Kien, B., et al. ABHD5 stimulates PNPLA1-mediated omega-O-acylceramide biosynthesis essential for a functional skin permeability barrier. J Lipid Res 59, 2360–2367 (2018).

24. Yang, A., Mottillo, E.P., Mladenovic-Lucas, L., Zhou, L. & Granneman, J.G. Dynamic interactions of ABHD5 with PNPLA3 regulate triacylglycerol metabolism in brown adipocytes. Nat Metab 1, 560–569 (2019).

25. Benner, C., et al. The transcriptional landscape of mouse beta cells compared to human beta cells reveals notable species differences in long non-coding RNA and protein-coding gene expression. BMC Genomics 15, 620 (2014).

26. Balboa, D., et al. Functional, metabolic and transcriptional maturation of human pancreatic islets derived from stem cells. Nat Biotechnol 40, 1042–1055 (2022).

27. Bronson, S.C., Shanmugam, A. & Mythili, C. Syndromic Conundrums in Diabetes: Seek and Ye Shall Find: The Dorfman-Chanarin Syndrome. Clin Diabetes 39, 117–120 (2021).

28. Schweiger, M., Lass, A., Zimmermann, R., Eichmann, T.O. & Zechner, R. Neutral lipid storage disease: genetic disorders caused by mutations in adipose triglyceride lipase/PNPLA2 or CGI-58/ABHD5. Am J Physiol Endocrinol Metab 297, E289–296 (2009).

29. Harata, M., et al. Delivery of shRNA via lentivirus in human pseudoislets provides a model to test dynamic regulation of insulin secretion and gene function in human islets. Physiol Rep 6, e13907 (2018).

30. Mishra, A., et al. Perilipin 2 downregulation in beta cells impairs insulin secretion under nutritional stress and damages mitochondria. JCI Insight 6(2021).

31. Brennecke, B.R., et al. Utilization of commercial collagens for preparing well-differentiated human beta cells for confocal microscopy. Front Endocrinol (Lausanne) 14, 1187216 (2023).

32. Trevino, M.B., et al. Perilipin 5 regulates islet lipid metabolism and insulin secretion in a cAMP-dependent manner: implication of its role in the postprandial insulin secretion. Diabetes 64, 1299–1310 (2015).

