## Supplementary Figure 1 for "ALPHA/BETA HYDROLASE DOMAIN CONTAINING 5 SUPPORTS GLUCOSE-STIMULATED-LIPOLYSIS AND INSULIN SECRETION IN PANCREATIC BETA CELLS"

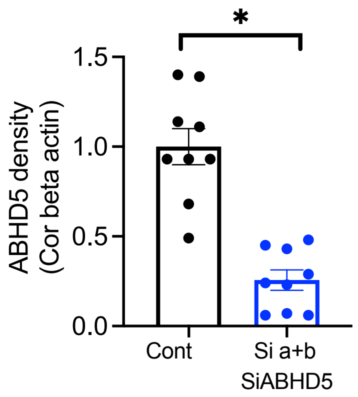


Supplementary Figure 1: INS-1 cells were transfected with non-targeting SiRNA (Cont) or the combination of two SiRNA against ABHD5 (Si a+b). Western blot of ABHD5 and beta-actin was performed and band intensity of ABHD5 corrected for beta-actin was expressed taking average value for Cont in each experiment as 1. n= 9. Mean ± sem. *; $p<0.05$ by Student’s t test.
